# Natural variations in maternal behaviour shape trait anxiety and hippocampal neurogenesis in offspring

**DOI:** 10.1101/2024.09.28.615571

**Authors:** Fabio Grieco, Pauline van Gelderen, Thomas Larrieu, Nicolas Toni

## Abstract

Understanding the early origins of emotional traits and baseline anxiety is crucial for the development of personalized medicine in mood disorders. While previous research primarily focused on extreme conditions like chronic maternal deprivation, in the present study, we investigated how natural variations in maternal care influence anxiety-related behaviour and neurogenesis in C57BL/6J offspring in a longitudinal manner. We observed that mothers engaging in low maternal care (LMC) displayed lower adult neurogenesis in both the olfactory bulb and the dentate gyrus of the hippocampus compared to high maternal care (HMC) mothers. We then observed that LMC-reared pups exhibited increased anxiety-related behaviour at postnatal day (PND) 5, 9, and 22. Furthermore, maternal behaviour induced the development of emotional individuality at early stage. This mood-related phenotype in LMC-reared offspring was associated with decreased neurogenesis after weaning at PND24. In another group of litters, we further examined neurogenesis at an earlier age (PND9) and already found a reduction in the population of adult neural progenitor cells and cell proliferation in the subgranular zone of the dentate gyrus of LMC-reared pups. These results highlight that natural variations in early life experiences such as maternal care, shape long-term brain plasticity and behaviour in offspring. This underscores the relevance of maternal care and adult neurogenesis in shaping personality-like traits related to mood disorders.

## Introduction

Despite sharing the same genetic background and controlled environmental conditions typically used in laboratory settings, C57BL/6J mice exhibit remarkable interindividual differences in anxiety levels and adult hippocampal neurogenesis^1^. These variations raise important questions about the underlying mechanisms driving such differences under physiological conditions that can subsequently lead to stress vulnerability^2^. While genetic homogeneity and consistent housing are expected to minimise behavioural and neurobiological variability, these findings suggest that additional factors such as early-life experiences, or subtle environmental influences, also referred to as the non-shared environment^3^ play significant roles in shaping individual traits in inbred mice. Among these factors, we recently found that natural variations in adult neurogenesis and anxiety are key determinants in social rank, where dominant individuals display heightened trait anxiety and greater stress susceptibility compared to their subordinate counterparts^4–6^. However, to study the emergence of emotional individuality, one of the earliest phenomena that might influencing these traits is maternal care.

In all mammals, maternal behaviour, usually emerging at or close to parturition is a cornerstone of both physical and brain development, profoundly influencing the well-being of offspring^7^. Human studies have highlighted the critical role of early maternal care and maternal bonding on a child’s emotional development and cognitive functioning: For instance, children who experience consistent and nurturing maternal care tend to develop secure attachments, exhibit lower levels of stress, and perform better in cognitive tasks^8,9^. Conversely, adverse childhood experiences, including neglect, are linked to higher risks of developing neuropsychiatric disorders^10^ and cognitive impairments^11^ later in life. Animal studies have long suggested that the quality and type of maternal care can affect the emotional behaviour of offspring, potentially predisposing them to anxiety-, depression-related behaviour, or resilience in the face of stress. Adult offspring of mothers engaging in high maternal care during the first postnatal week show less anxiety-like behaviour and reduced hypothalamic– pituitary–adrenal (HPA) stress responses^12–14^. A wide range of behavioural changes during both pregnancy and postpartum are present in all females. In rodents, these changes include enhanced nesting activity before birth, known as “preparatory nesting,” to create a shelter for the upcoming litter^15,16^. After birth, the female shows a very immediate interest in the newborn reflected by key behavioural features such as liking, retrieval, and gathering^17^. However, the most important and common pattern of maternal behaviour in mammals, including rodents is nursing. Once the dam has retrieved and gathered the pups together, it spends most of its time in the nest in a nursing posture^18^. This nursing behaviour is more prevalent during the light phase, coinciding with periods of increased inactivity^19^. At birth, newborn mice are naked, blind, deaf, and they depend entirely on their mother for warmth, nourishment, and shelter. For their survival, this implies that they remain close to each other and the mother in the nest during early development. It is not surprising then that natural variations in maternal care, even seen in inbred strains such as C57BL/6 mice, can lead to diverse behavioural and physiological outcomes in the adult offspring^12,20^. These emotional trajectories indicate a long-term influence of early life experiences such as maternal care.

A hallmark of mood-related behaviour is the modulation of adult neurogenesis (AN), which primarily occurs in the subventricular zone (SVZ)^21^ and the subgranular zone (SGZ) of the dentate gyrus (DG)^22^. In the SVZ, newly generated neurons migrate via the rostral migratory stream (RMS) to the olfactory bulb (OB), where they mature and enhance olfactory function crucial for odor detection and mother-pup interactions^23–25^. Within the hippocampus, these neurons integrate into the network^26^, playing a vital role in modulating hippocampal activity and regulating stress responses^27–31^. These neurons are highly plastic and are shaped by various environmental factors such as physical activity, aging, nutrition, disease, and stress^32–38^. Consequently, AN serves as both a key regulator of both olfactory bulb and hippocampal functions and a biomarker for overall brain health. Artificial or conditioned low maternal care modelised by chronic maternal separation (MS) protocols indicate that prolonged MS inhibits AN and emotional behaviour in adult offspring^39^. However, while MS or deprivation studies reflect the importance of mother–infant interaction for hippocampal neuronal development in rodents^39–43^, they do not directly address the issue of whether under normal conditions, natural variations in maternal care influence baseline offspring AN along with anxiety-related traits at early stage of life. In addition, little is known about the putative link between natural variations observed in maternal care and baseline adult neurogenesis in the mother itself.

## Results

### Mothers with spontaneously low maternal care display less neurogenesis in the olfactory bulb and hippocampus

Rodents exhibit natural variations in maternal care^12,20^. As to whether these inter-individual differences are associated with differences in the levels of adult neurogenesis is not known. We observed and scored the behaviour of dams with their pups during the first two weeks post-parturition, monitoring individual variations in maternal parameters including arched-back nursing (ABN) and licking-grooming (LG) of pups (Fig. 1A)^12^. In addition to scoring these standard parameters (Fig. 1B), we also used occurrence in the nest (ON) as a simple method to assess maternal care. By segregating mice into two groups based on the mean total ON, we identified two distinct maternal profiles: high maternal care (HMC) and low maternal care (LMC) (Fig. 1C). Pup retrieval test and ABN/LG behaviours are well-established gold standard to assess maternal care^44–46^. Importantly, our analysis revealed a strong correlation between the percentage of ON and established maternal behaviours such as ABN and LG (R^2^ = 0.842; P < 0.001). We further validated this method by showing that mothers exhibiting low levels of maternal care demonstrated significantly higher latency in retrieving pups within the home cage at postnatal day 5 (PND5), indicating reduced maternal responsiveness to separation (Fig. 1D). This LMC profile was also evidenced by a significant reduction in the proportion of pups with visible milk spots at PND5 and PND6, highlighting compromised active nursing behaviour (Fig. 1E). Finally, principal component analysis (PCA) revealed a clear separation between the LMC and HMC, with no overlap in behavioural patterns, supporting the validity of our classification system using ON (Fig. 1F, G). This finding strongly indicates that ON is a reliable parameter to simplify the evaluation of maternal behaviour. Indeed, assessing the time spent in the nest offers a more straightforward, reliable, and reproducible measure of ‘maternal behaviour’ compared to individually monitoring passive nursing, arched-back nursing, pup contact, and licking-grooming, as it integrates all these behaviours, which predominantly occur within the nest environment. Importantly, neither prenatal preparatory nesting behaviour (Fig. 1H) nor baseline anxiety levels (Fig. 1I) of the mothers predicted maternal care quality, indicating these factors might be independent of postpartum maternal behaviour.

**Figure 1.**
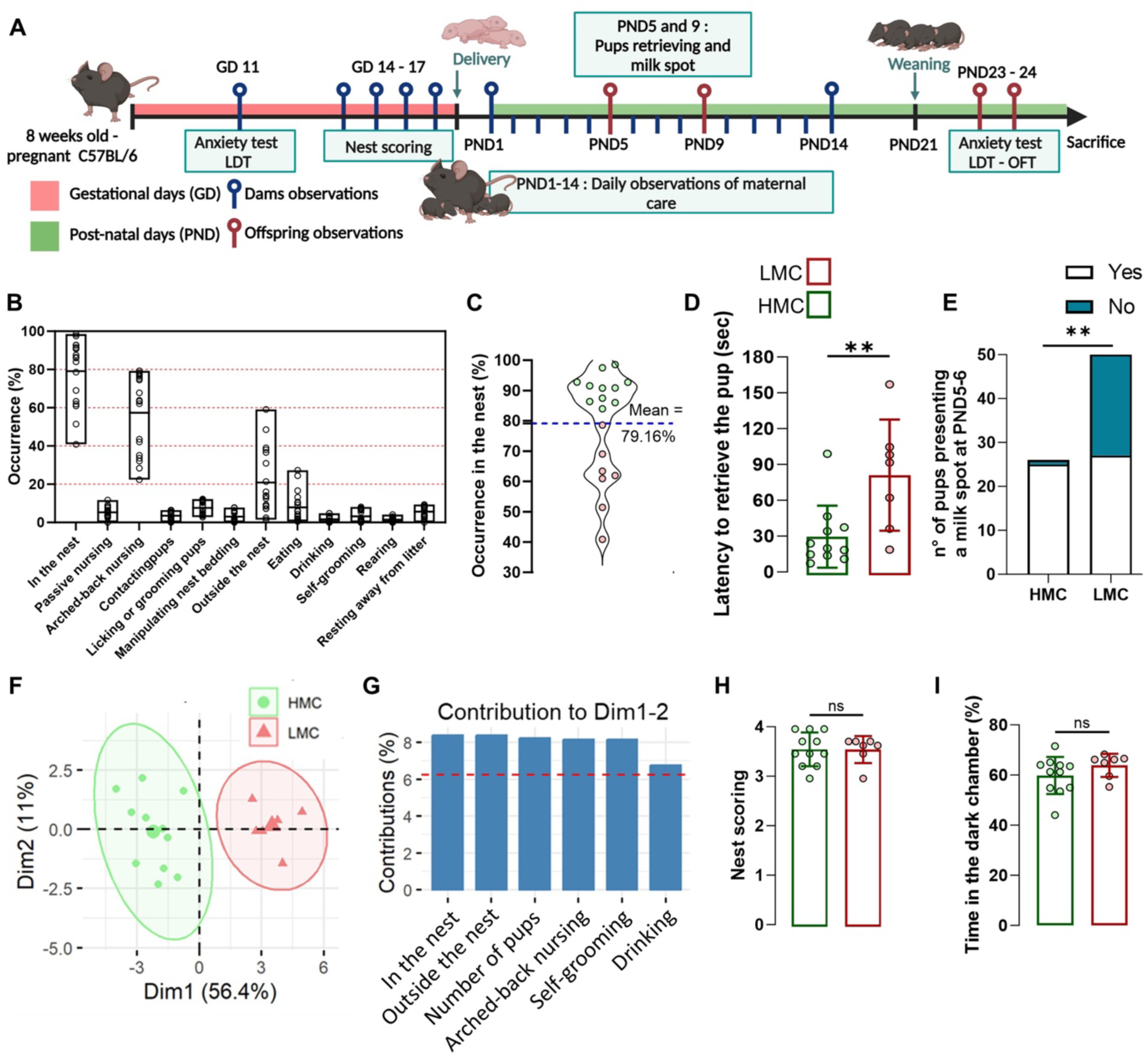
Occurrence in the nest assesses natural variation of maternal care: **A.** Schematic illustration of the experimental design. **B.** Histograms of the percentage of daily observations of maternal behaviours of low (LMC) versus high maternal care (HMC) mothers. **C.** Violin plot showing the distribution and the mean of occurrence in the nest used to discriminate LMC from HMC mothers. **D.** Latency to retrieve the pups at PND5 (t16=3.03, p = 0.008, unpaired t test, two-tailed, n = 11 and 7 per group). **E.** Number of pups presenting a milk spot at PND5-6 (Fisher’s exact test, two-sided, p = <0.0001). **F.** Samples representation using the principal component analysis (PCA) (n = 18). **G.** Relative contribution of the behavioural parameters to the dimensions 1 and 2 represented in the PCA. **H.** Nest quality score assessed at gestational days 14 and 15 resulting from the ability to make a nest (Mann-Whitney test, p = 0.993, two-tailed, n = 11 and 7 per group). **I.** Time spent in the dark chamber during a LDT at GD 11 (t16=1.28, p = 0.219, unpaired t-test, two-tailed, n = 11 and 7 per group). Histograms show average ± SEM, * p<0.05, ** p<0.01, ***p<0.001, ns = not significant.

At the cellular level, LMC mothers displayed reduced adult neurogenesis, as revealed by a reduction in the number of DG and OB cells expressing the immature neuronal marker, Doublecortin (DCX) (Fig. 2A-D). To further elucidate the relation between behaviour and neurogenesis, we conducted a correlation matrix encompassing various behavioural and cellular parameters (Fig. 2E). Interestingly, the latency to initiate pup retrieval negatively correlated with adult neurogenesis in the OB and DG of the mother while ABN/LG positively correlated with adult neurogenesis (Fig. 2E). In addition, the occurrence in the nest showed a positive correlation with adult neurogenesis in the OB and DG of the mother, suggesting that natural variations of adult neurogenesis in these brain regions under physiological conditions are intricately linked with interindividual differences in maternal behaviour. Strikingly, this correlation analysis revealed a significant association between ON, pup retrieval and ABN/LG parameters and litter size reflecting an important role of the number of pups per litter on the quality of maternal behaviour quality (Fig. 2E). Overall, our findings demonstrate that maternal behaviour quality can be distinctly classified based on nest occurrence, with a “bad mother” profile being closely associated with reduced adult neurogenesis in both the hippocampus and olfactory bulb.

**Figure 2.**
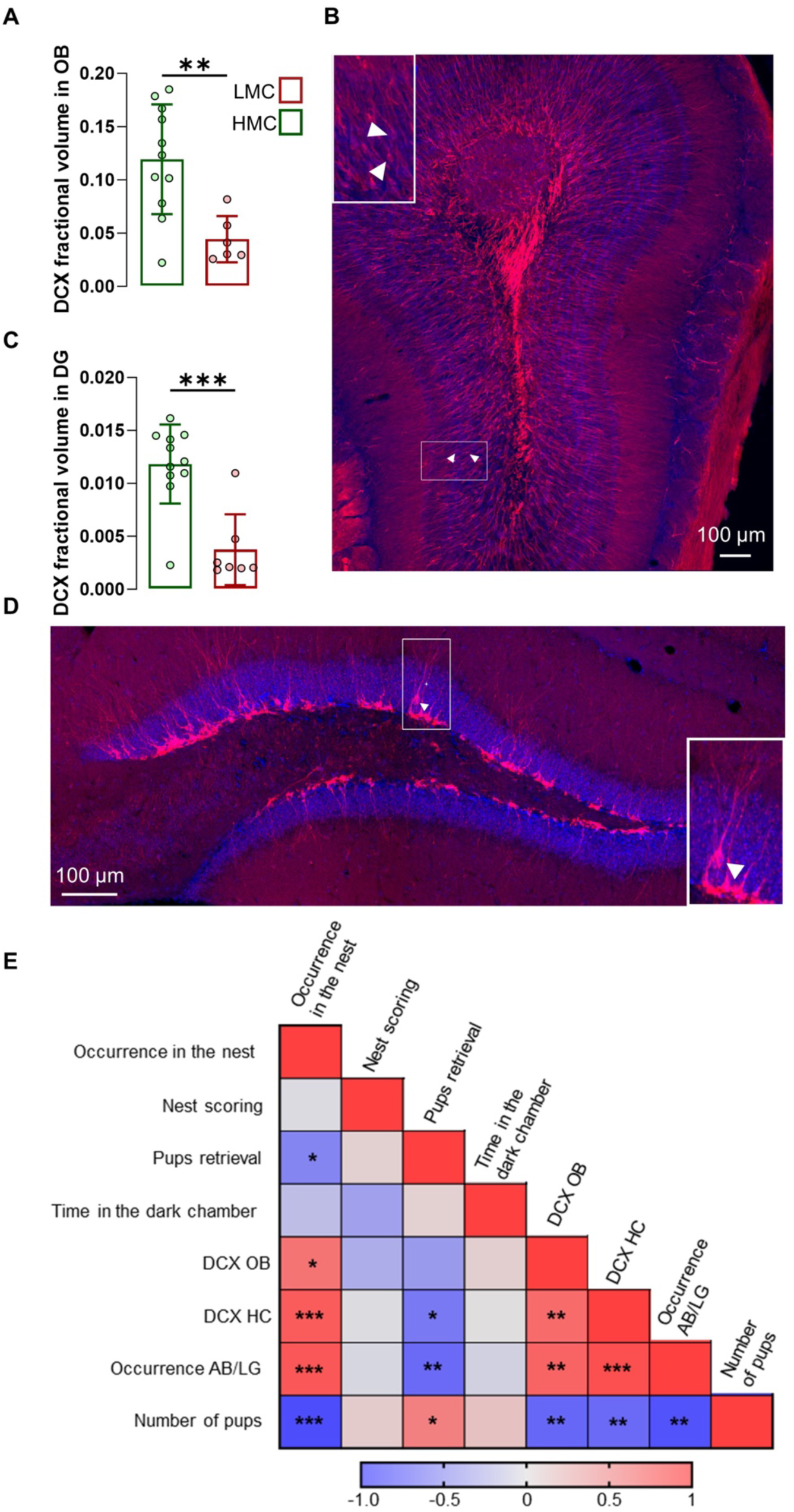
LMC mothers displayed decreased adult neurogenesis in the hippocampus and olfactory bulb. **A.** DCX fractional volume in the dentate gyrus of LMC versus HMC (t16=4.66, p = <0.001, unpaired t test, two-tailed, n = 11 and 7 per group). **B.** DCX fractional volume in the olfactory bulb (t15=3.37, p = 0.004, unpaired t-test, two-tailed, n = 11 and 6 per group). **C, D.** Confocal maximal projections of the dentate gyrus and olfactory bulb, immunostained for DCX (red), and DAPI (blue). **E.** Heat map matrices representing Pearson’s correlation between maternal behaviour parameters, DCX fractional volume in OB and hippocampus, prenatal anxiety, and number of pups. Histograms show average ± SEM, * p<0.05, ** p<0.01, ***p<0.001, ns = not significant.

### Natural variations in maternal care influence pups’ anxious behaviour and emotional individuality

Ultrasonic vocalisations (USVs) are crucial for mother-pup communication, particularly in scenarios involving impaired social interactions, such as maternal deprivation^47,48^. Pup USVs have been considered a reliable indicator of anxiety or distress in response to maternal separation, a perspective supported by studies demonstrating that anxiolytic agents reduce USV emission^49,50,51^. Here we assessed the relationship between maternal care and anxiety-related behaviour of offspring using USVs. USVs emission in mouse pup peaks during the first postnatal week^52,53^. Hence, we measured USVs at PND5 and PND9 to observe developmental changes in vocalisation as the pups matured (Fig. 3A). When separated from their mothers, LMC male and female pups exhibited a heightened frequency (Fig. 3B) and duration (Fig. 3C) of vocalisations, compared to pups raised by HMC mothers. In pup mice, USVs can be divided into simple USVs which are characterized by single, short-duration, frequency-modulated calls and more elaborated calls such as ‘complex’, ‘two-components’, ‘composite’, ‘harmonic’ and ‘frequence steps’ that exhibit multiple frequency modulations, varied pitch, and intricate temporal patterns (Supplementary Fig 1D, E)^54^. At PND5, we found that although LMC and HMC pups did not differ in terms of simple calls (Fig. 3D), they did differ in terms of more elaborated calls reflected by an increased proportion of ‘complex’, ‘composite’ and ‘frequence steps’ syntax in LMC pups (Fig. 3E and Supplementary Fig. 1D). In addition, similar to a mouse model of autism ^55^, mean peak frequency was decreased (Fig. 3F) and amplitude was increased (Fig. 3G) in LMC-pups compared to HMC-pups. A similar but more pronounced pattern was observed at PND9 (Fig. 3H-M), suggesting that the influence of maternal care on vocalisation is established early in development and persists as the pups mature. Finally, LMC mice showed a significantly greater variance in most of the USV parameters we measured at both PND5 and PND9, suggesting that low maternal behaviour does stimulate the development of interindividual differences in isolation-induced vocalisation emissions in pups at this early stage. To strengthen this observation with more sample size, we pooled data from 2 independent experiments at PND9 (Supplementary Fig. 2) and found that LMC mice showed a significantly greater variance in all the USV parameters we measured except for the proportion of complex calls, confirming that low maternal care stimulates the development of emotional individuality in isolation-induced anxiety-related behaviour in pups at early stage. Importantly, our observations at PND5 and PND9 revealed similar profile between LMC and HMC pups in terms of body weight, righting reflex, and cliff avoidance tests, suggesting that there is no developmental delay in LMC as compared to HMC pups (Supplementary Fig. 1A-C). Taken together, the analysis of USV indicates that low maternal care elevates the probability of vocalisations in both sexes at PND5 and PND9 during acute maternal separation and reveals the individualisation of behaviour.

**Figure 3.**
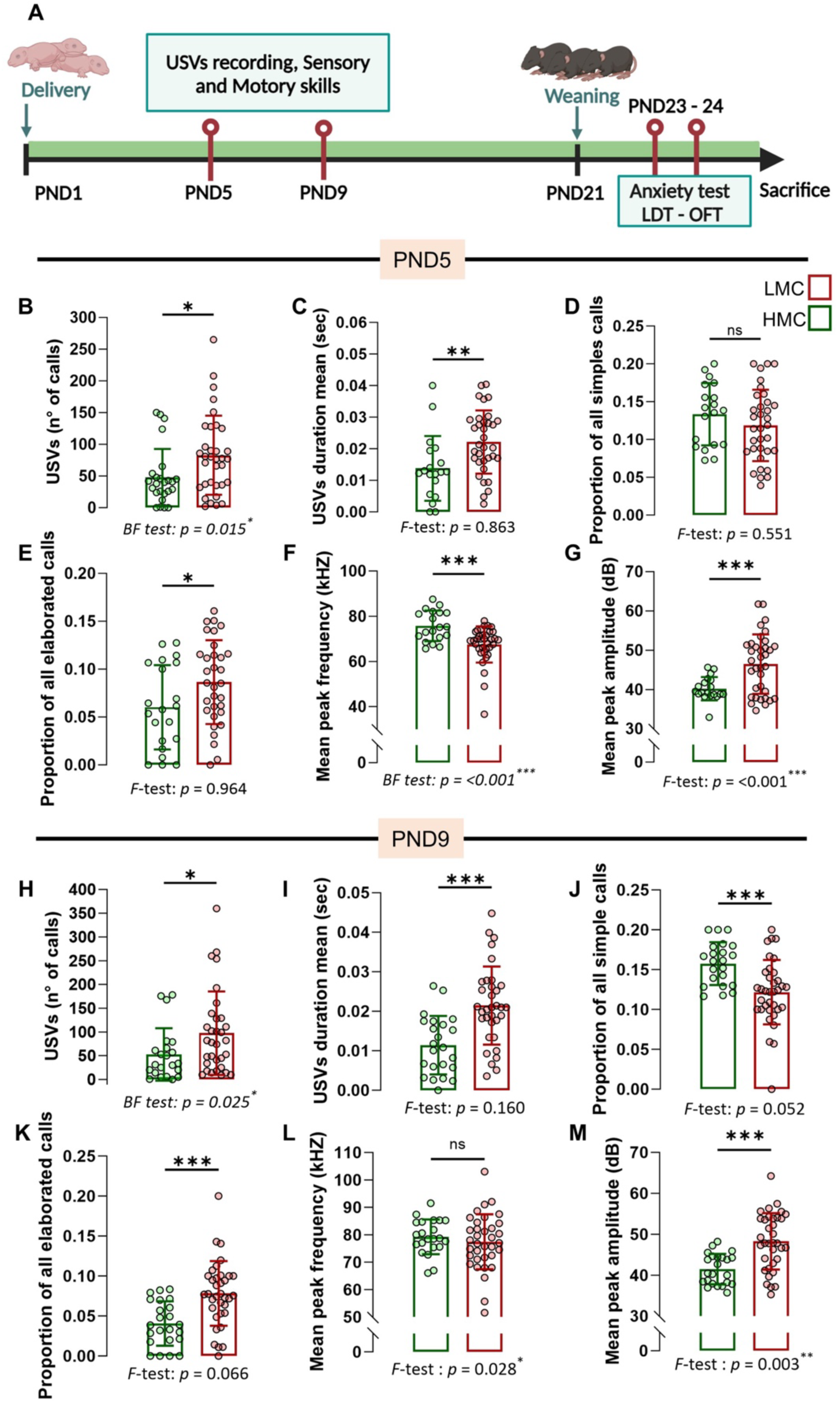
LMC-reared offspring displayed anxiety-related behaviour assessed by USV recordings. **A.** Experimental design for anxiety-related behaviour of offspring reared by LMC and HMC mothers at PND5, PND9, and PND23. **B.** Total numbers of calls upon acute maternal separation (Mann-Whitney test, p = 0.021, two-tailed, n = 25 and 34 per group). **C.** Mean duration of the calls (t51=2.89, p = 0.006, unpaired t test, two-tailed, n = 19 and 34 per group). **D.** Proportion of all simple calls (t51=1.14, p = 0.256, unpaired t-test, two-tailed, n = 19 and 34 per group). **E.** Proportion of all elaborated calls (t51=2.13, p = 0.038, unpaired t test, two-tailed, n = 21 and 32 per group). **F.** Mean peak frequency (Mann-Whitney test, p = <0.001, two-tailed, n = 19 and 35 per group). **G.** Mean peak amplitude (t48.62=4.32, p = <0.001, Welch’s test, two-tailed, n =19 and 35 per group). **H**. Total number of calls (Mann-Whitney test, p = 0.033, two-tailed, n = 22 and 32 per group). **I.** Mean duration of calls (t55=4.14, p = <0.001, unpaired t-test, two-tailed, n = 23 and 34 per group). **J.** Proportion of all simple calls (t55=3.67, p = <0.001, unpaired t-test, two-tailed, n = 22 and 35 per group). **K.** Proportion of all elaborated calls (t56=3.90, p = <0.001, unpaired t-test, two-tailed, n = 23 and 35 per group). **L.** Mean peak frequency (t54=0.84, p = 0.402, Welch’s test, two-tailed, n = 22 and 35 per group). **M.** Mean peak amplitude (t54=4.86, p = <0.001, Welch’s test, two-tailed, n 23 and 35 per group). Histograms show average ± SD, *p<0.05, ** p<0.01, ***p<0.001, ns = not significant.

### LMC increases trait anxiety in weaned offspring but does not affect individuality

To investigate the potential influence of natural variations in maternal care on trait anxiety in offspring after weaning, the same mice were evaluated on a Light-Dark Test (LDT) and an Open Field Test (OFT), at PND 22-23 (Fig. 4A-C). We found that LMC offspring exhibited increased anxiety-related behaviour as compared to HMC pups, revealed by increased time spent in the dark chamber of the LDT (Fig. 4A) and in thigmotaxis during an OFT (Fig. 4B). A composite anxiety score, calculated from normalised values of both tests, revealed higher anxiety levels in the LMC group (Fig. 4C). Importantly, there was no significant difference in the distance travelled between the groups in either the LDT or the OFT, indicating that variations in anxiety-related behaviour were not confounded by differences in overall locomotor activity (Supplementary Fig. 3A, B). Finally, the groups did not differ in the variances in both le LDT and the OFT, suggesting that LMC did not trigger the development of interindividual differences in anxiety-related behaviour at this age. Overall, these findings support the hypothesis that natural variations in maternal care quality have a significant impact on the development of trait anxiety in juvenile mice, highlighting the critical role of early life experiences in shaping emotional behaviour.

**Figure 4.**
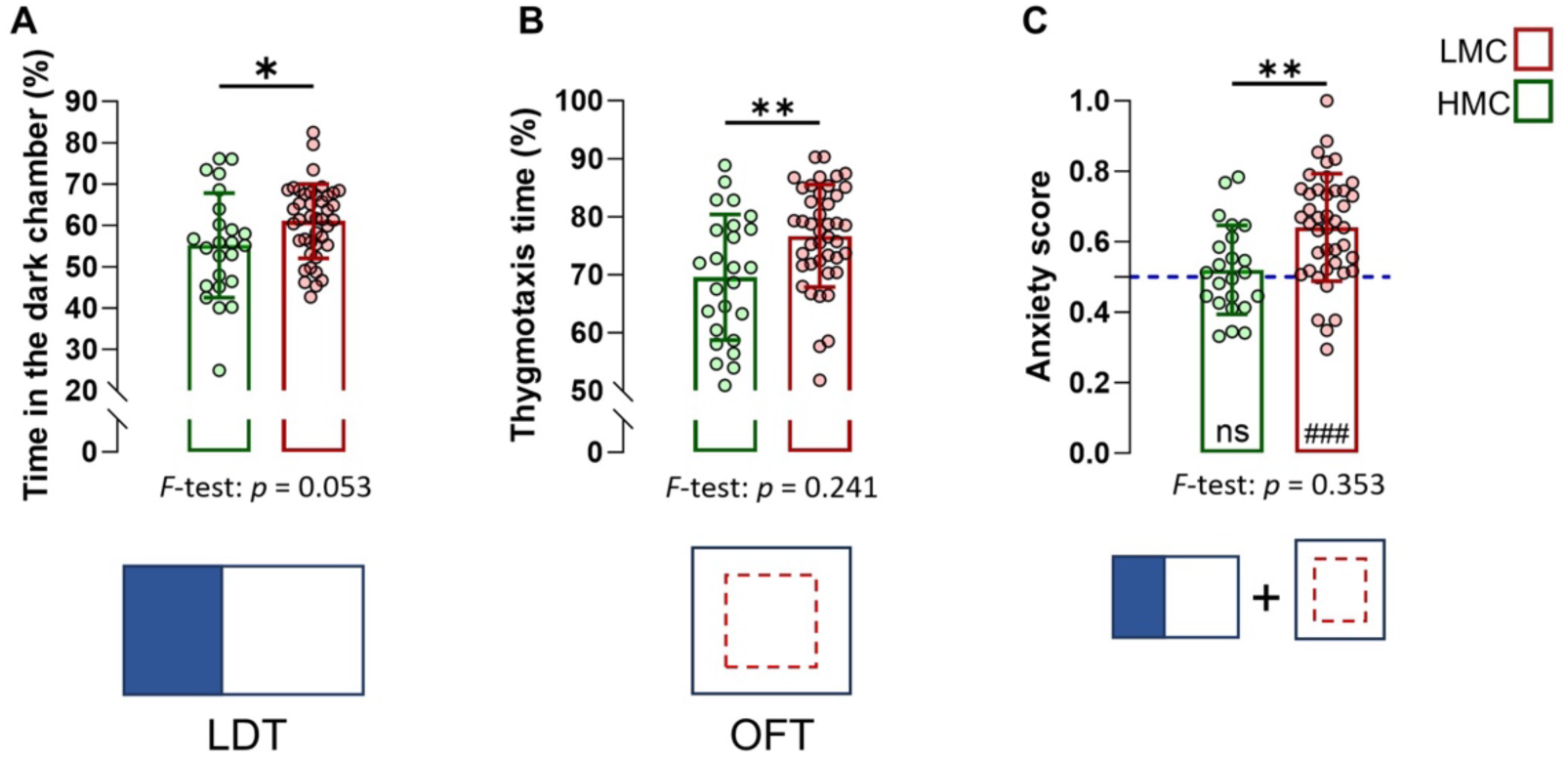
LMC-reared offspring displayed increased anxiety-related behaviour at weaning. **A.** Time spent in the dark chamber during a light-dark test at PND22 (t64=2.18, p = 0.032, unpaired t-test, two-tailed, n = 24 and 42 per group). **B.** Time spent in thigmotaxis during the open field at PND23 (t65=2.93, p = 0.005, unpaired t-test, two-tailed, n = 25 and 42 per group). **C.** Anxiety score derived from normalized data from LDT and OFT (t54=3.22, p = 0.002, unpaired t-test, two-tailed, n = 23 and 42; t41=5.99, p = <0.0001, one sample t-test, two-tailed, n = 42). Histograms show average ± SD, * p<0.05, ** p<0.01, ***p<0.001, ns = not significant. Comparison between the group mean and the hypothetical value of 0.5, to assess anxiety withing each group are shown within each histogram bar. One-sample t-test: #p<0.05, ##p<0.01, ###p<0.001, ns = not significant.

### LMC decreases levels of neurogenesis in weaned offspring

Natural variations in anxiety-related behaviour under baseline conditions relate to variations in adult neurogenesis^1,6^. To investigate whether the observed behavioural variability in the offspring was associated with differences in adult neurogenesis in the DG of the hippocampus, we assessed the proliferation of precursor cells by staining mouse brain sections with the cell proliferation marker Ki67. Additionally, we identified newborn neurons undergoing maturation by the presence of doublecortin (DCX). Lower numbers of proliferating cells in the subgranular zone of the DG (Fig. 5A, C) and DCX fractional volume (Fig. 5B, D) was observed in LMC-compared to HMC-reared juvenile mice. These findings suggest that reduced maternal care adversely affects neurogenesis in weaned offspring, potentially underlying the observed behavioural differences, and highlight the critical role of early-life experiences in shaping brain development and function in offspring. To strengthen this idea, we conducted a correlation matrix encompassing various behavioural and cellular parameters of the offspring at different ages and maternal care (Fig. 5E). Correlation analyses further revealed that adult neurogenesis in the hippocampus of weaned offspring negatively correlated with their anxiety score as previously observed in adult^1^. Strikingly, occurrence in the nest during maternal care observation predicts the levels of adult neurogenesis in juvenile offspring as revealed by the positive correlation between ON parameter and both Ki67 and DCX positive cells in the dentate gyrus (Fig. 5E). We found no significant difference in the variance of markers of adult neurogenesis between LMC and HMC juveniles.

**Figure 5.**
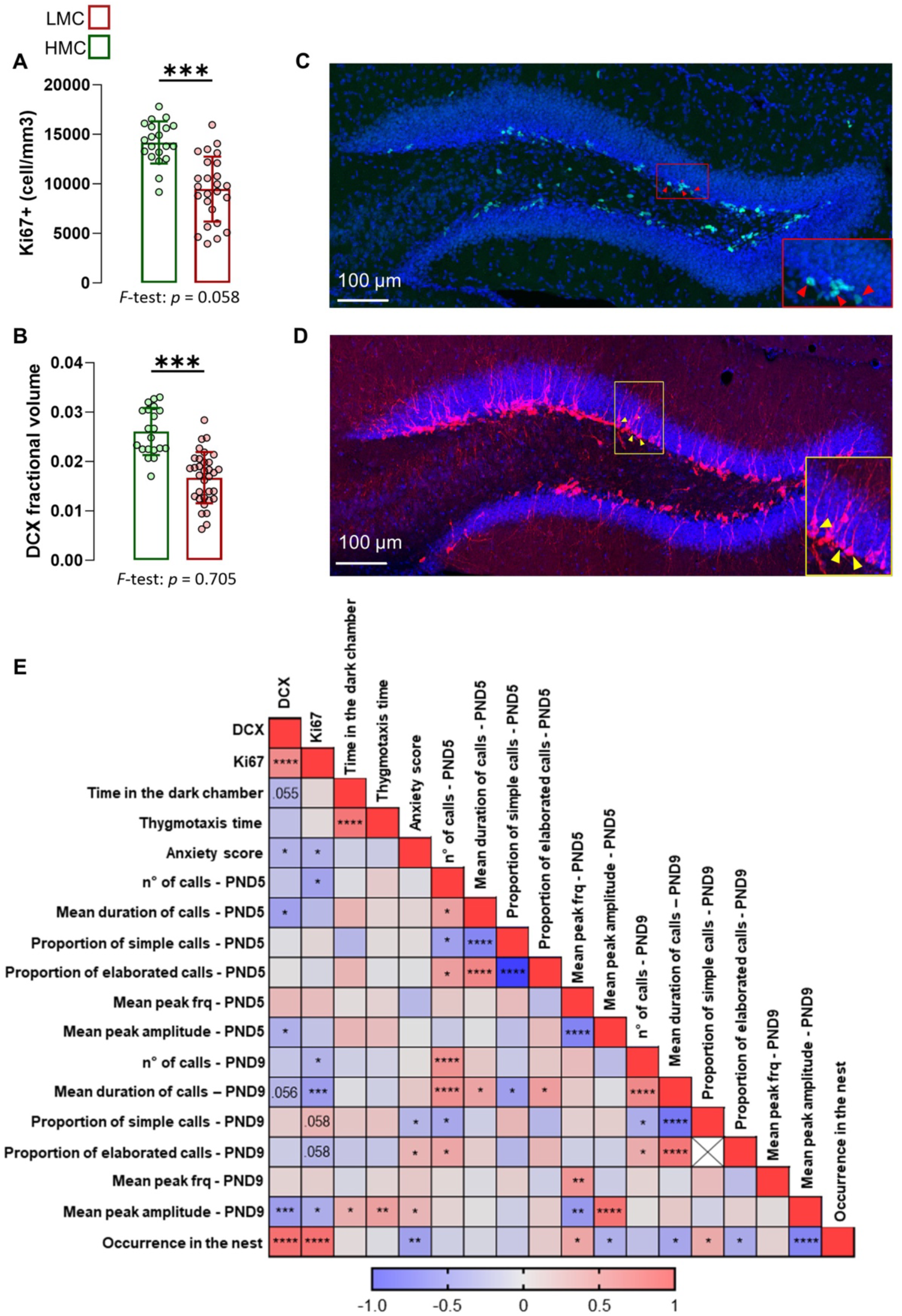
Juvenile LMC-reared offspring displayed decreased hippocampal adult neurogenesis. **A.** Quantification of Ki67 positive cells in the dentate gyrus of the hippocampus of LMC- and HMC-reared offspring (t43=5.55, p = <0.001, unpaired t test, two-tailed, n = 25 and 20 per group). **B.** DCX fractional volume in the dentate gyrus of the hippocampus of LMC- and HMC-reared offspring (t51=6.55, p = <0.001, unpaired t-test, two-tailed, n = 20 and 33 per group). **C, D.** Representative confocal microscopy image of the dentate gyrus immunostained for Ki67 (**C.** red cells highlighted with yellow arrows), and DCX (**D.** white cells highlighted with yellow arrows) and DAPI (**C and D.** blue). **E.** Heat map representing Pearson’s correlation matrices between selected USVs features, adult neural progenitor cell proliferation and immature neurons in the dentate gyrus, juvenile anxiety and the time spent in the nest by the mother. Histograms show average ± SD, * p<0.05, ** p<0.01, ***p<0.001, ns = not significant. Inset scale bars: 10μm.

### LMC decreases levels of adult neural progenitors and their proliferation at PND9

Building on our prior findings where LMC-reared pups at PND9 exhibited increased USV calls and a reduction in adult neurogenesis at PND24, we sought to investigate whether these differences in neurogenesis could emerge at an earlier age, i.e. PND9. To address this, we conducted a second longitudinal experiment where we first observed and scored maternal behaviour from PND1 to PND9 as described above and segregated mothers into HMC and LMC groups based on the mean total ON, (Supplementary Fig. 4). Importantly, the analysis of maternal care in experiment 1 revealed a strong correlation between the percentage of ON assessed during the first two weeks post-parturition and the first week (Supplementary Fig. 4A), indicating that 9 days of observation are sufficient to classify maternal care. Notably, in this second cohort of mice, we were able to reproduce USV phenotypes in LMC-reared pups at PND9 strengthening our initial observations (Fig. 6A-G). Most importantly, LMC pups displayed a significant decrease in the numbers of proliferating cells in the subgranular zone of the DG (Fig. 6H, J), HopX-expressing stem cells (Fig. 6H, I), and proportion of HopX^+^ stem cell that proliferated (Fig. 6K) as compared to HMC-reared pups. At this developmental stage, we found statistical differences in the variances of Ki67 positive cells between groups (Fig. 6J). These findings suggest that low maternal care induces detrimental effects on neurogenesis at early stage of postnatal development, potentially underlying the observed behavioural differences with moderate impact on individualisation in the HMC-reared pups. In addition, correlation analyses further revealed the close relationship between proliferating cells in the dentate gyrus of PND9 pups and USV parameters (Fig. 6L). Finally, ON parameter during maternal care observations predicts the levels of adult neural progenitor cell proliferation in the dentate gyrus of the hippocampus of PND9 pups (Fig. 6L) revealing a crucial role of maternal care quality in brain plasticity in offspring at an early stage of development.

**Figure 6.**
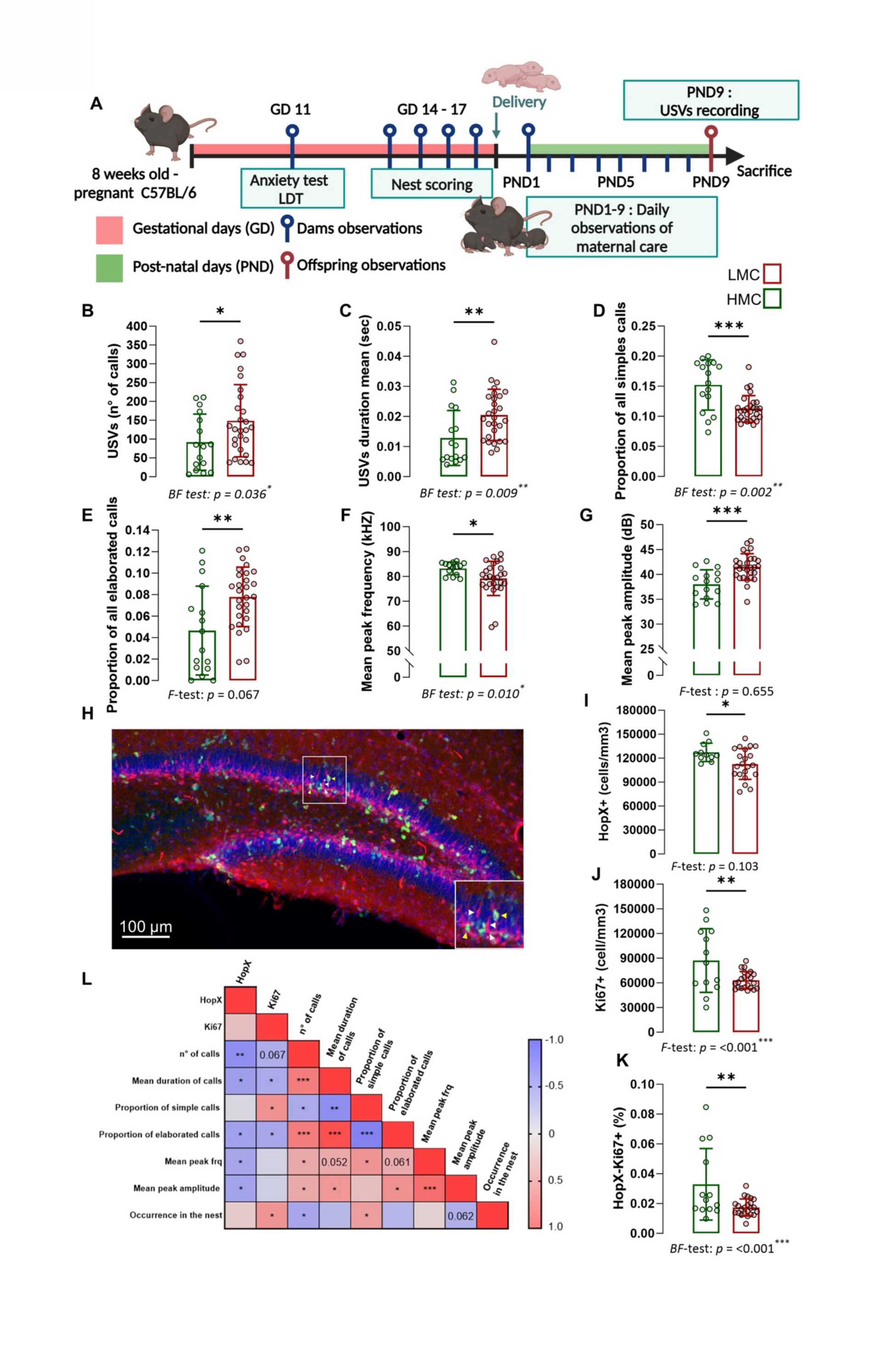
LMC-reared offspring displayed decreased hippocampal neurogenesis at PND9. **A.** Schematic representation of experimental design. **B.** Total numbers of calls upon acute maternal separation (Mann-Whitney test, p = 0.039, two-tailed, n = 16 and 27 per group). **C.** Mean duration of the calls (Mann-Whitney test, p = 0.004, two-tailed, n 16 and 28 per group). **D.** Proportion of all simple calls (Mann-Whitney, p = 0.003, two-tailed, n = 16 and 28 per group). **E.** Proportion of all elaborated calls (t42=3.01, p = 0.004, unpaired t test, two-tailed, n = 16 and 28 per group) **F.** Mean peak frequency (Mann-Whitney test, p = 0.036, two-tailed, n = 14 and 27 per group). **G.** Mean peak amplitude (t41=3.95, p = <0.001, unpaired t-test, two-tailed, n =15 and 28 per group). **H.** Confocal maximal projection of the dentate gyrus immunostained for Ki67 (in red, highlighted with yellow arrows) and HopX (in green, highlighted with yellow arrows). **I.** Quantification of HopX positive cells (t30=2.31, p = 0.027, unpaired t test, two-tailed, n = 11 and 21 per group). **J.** Quantification of Ki67 positive cells (t13.05=2.18, p = 0.048, Welch’s test, two-tailed, n = 13 and 22). **K.** Percentage of HopX^+^-Ki67^+^ cells over the total amount of HopX^+^ cells (t31=2.35, p = 0.025, unpaired t-test, two-tailed, n = 12 and 21 per group). **L.** Heat map shows Pearson’s correlation between selected USV features, cellular proliferation and neuronal progenitors in the dentate gyrus. Histograms show average ± SD, * p<0.05, ** p<0.01, ***p<0.001, ns = not significant. Inset scale bars: 10μm.

## Discussion

Epidemiological and clinical research indicate that adverse early-life experiences represent significant risk factors for the development of psychopathologies in offspring^10^. Using a longitudinal behavioural approach for assessing maternal care, we found that natural variations in maternal care quality are associated with interindividual differences in neurogenesis in the olfactory bulb (OB) and the dentate gyrus (DG) of dams which, in turn, significantly influence anxiety traits and neurogenesis in their offspring during early developmental stages (PND9 and PND23).

We found that maternal behaviour is not uniform even among the same inbred C57Bl/6J strain and that mothers can be segregated into low maternal care (LMC) and high maternal care (HMC) which is consistent with previous studies showing individual variations in maternal styles after 2-week monitoring of dams with their pups^12,19^. Here we combined established gold standard parameters with a new and simple method that evaluates maternal care based on the occurrence in the nest (ON)^56^. This approach revealed a strong correlation between ON and traditional maternal behaviours such as arched-back nursing (ABN) and licking-grooming (LG)^12^, validating ON as a reliable proxy for comprehensive maternal behaviour assessment. Moreover, this method circumvents the need for separate monitoring of passive nursing, ABN, pup contact, and LG, thereby enhancing observational efficiency and consistency. Further validation of the ON parameter revealed that LMC mothers exhibited significantly higher latencies in pup retrieval within the home cage at postnatal day 5 (PND5), indicative of reduced maternal responsiveness to pup separation as observed after gestational stress^44^. To strengthen the utility of ON method, the LMC profile was also characterized by a substantial reduction in the proportion of pups with visible milk spots at PND5 and PND6, indicating that nursing behaviour was severely impaired^23^.

As olfaction is essential in mice to maintain appropriate maternal behaviour^57^, and previous studies linked motherhood to increased neurogenesis in the OB^58,59^, we compared neurogenesis in the OB and the DG of LMC and HMC mothers. We found that LMC mothers, as compared to HMC mothers, exhibited a decrease in adult neurogenesis, indicated by a reduced number of cells expressing the immature neuronal marker Doublecortin (DCX) in both the DG and the OB. This result is in line with previous studies reporting that normal expression of postpartum maternal behaviour requires increased adult neurogenesis in the SVZ of pregnant female mice^58,59^. Interestingly, nulliparous female rats exposed to pups that display maternal care show more proliferating cells in the SVZ compared to females never exposed to pups or females who do not display any maternal care^60^. This suggests that the act of providing maternal care itself may be an important modulator of neurogenesis, regardless of pregnancy or actual motherhood.

To investigate the impact of interindividual differences in maternal care on pup behaviour at an early stage of life, we measured USV emissions at PND5 and PND9. During the perinatal and early postnatal periods of rodent development, USVs constitute the primary acoustic signaling mechanism that serves as a crucial ethological biomarker to assess the emotional state of the pups^61^. Transient maternal separation is a well-known paradigm for eliciting USVs in rodent offspring. This acute separation-induced USVs increases during the first week of pup life, reaching a peak around PND7, and decreasing until PND14^53^. Here, we observed that LMC resulted in higher USV emissions in pups, reflected by an increase in the number and duration of calls. The increase in vocalisations in LMC-reared pups suggests a unique emotional response to separation when compared to HMC-reared pups^48^, as supported by studies indicating increased levels of anxiety-related behaviour in adult offspring mice raised by ‘bad’ mothers^12,20^. Further supporting this idea, anxiolytic compounds suppress calling behaviour without affecting motor function in rat pups^62^ and selective serotonin reuptake inhibitors abolish the emission of separation-induced USVs^63^. We observed that deficiency in maternal behaviour also resulted in differences in transitioning between simple and more elaborated calls in pups. Hence, we found significant differences in the syntactic use of the ten different types of USVs. For instance, LMC pups exhibited a decrease in the proportion of ‘short’ and ‘down’ calls referred to as ‘simple calls’ while ‘complex’, ‘two-components’ and ‘composite’ calls referred to as ‘elaborated calls’ were increased. Our data illustrate the impact of maternal care quality on communication patterns during early postnatal development. Importantly, developmental motor and sensory skills were similar between groups, excluding general developmental delays that could underlie the observed differences in USVs.

We then investigated the impact of maternal care on anxiety-related behaviour in offspring after weaning. We confirmed that natural variations in maternal behaviour significantly influences anxiety-related behaviours in offspring using gold standard behavioural tests^12^. However, prior to our study, there had been no research into this emotional behaviour at such an early age, mainly PND22. Offspring from LMC mothers exhibited heightened anxiety, as evidenced by an increased time spent in the dark chamber of a light-dark box and in thigmotaxis behaviour during an open field test. This increased anxiety was not attributable to differences in general locomotor activity, suggesting a direct link between maternal care and the development of anxiety traits that are already evident in juveniles. Importantly, most of our understanding regarding the influence of maternal care on anxiety in offspring has come from studies using extreme experimental interventions such as chronic maternal deprivation^64,65^. In contrast, our current findings highlight that even subtle, naturally occurring variations in maternal care can lead to profound alterations in pups’ behaviour.

Emotional individualisation refers to the development of unique emotional traits and responses in individuals, shaped by a combination of genetic, environmental, and neurobiological factors. It suggests that even among genetically similar individuals, like inbred C57BL/6J mice, subtle early-life experiences such as maternal care or environmental enrichment can create significant variability in emotional behaviour^66^. We identified LMC behaviour as a valuable model for exploring the development of emotional individuality in offspring. Indeed, our longitudinal analysis reveals that LMC affects both the mean and variance of early behavioural traits. Notably, anxiety-related behaviours using USVs at early stages (PND5 and PND9) showed increased variability among individuals raised by bad mothers while anxiety traits were uniformly affected after weaning. These findings suggest that natural variations in maternal care can accentuate differences within a population, contributing to the emergence of unique emotional profiles at early stages of pup’s development as PND5 and PND9. This is in line with other study showing that environmental enrichment in rats, influences neurogenesis and behavioural outcomes and modulates inter-individual variability. Surprisingly, although it has been proposed that individualisation arises from the progressively differential response to non-shared environmental components^66^, our findings suggest that even in standard housing conditions, i.e. in absence of environmental enrichment, LMC can still drive emotional individuality. This observation highlights the potent influence of early maternal care on shaping emotional diversity in the pups and raises the possibility of a trans-generational amplification of individuality mediated by adult neurogenesis and maternal care.

Mood-related behaviours have long been linked to alterations in adult neurogenesis^67^. Here, our results reveal a robust negative correlation between individual trait anxiety and adult neurogenesis in juvenile offspring at PND24. Specifically, we observed reduced cell proliferation and immature neurons in the DG of the hippocampus in juvenile that were reared by LMC mother compared to HMC pups. This alteration in neurogenesis indicates that 21 days of maternal care (from PND1 to PND21) already shape brain of the juvenile offspring at a cellular level.

Strikingly, we observed that this alteration in adult neurogenesis of LMC-reared pups was already evident at PND9 after only 1 week of maternal care. However, further investigations are warranted to delve into the underlying involvement of genomic events, such as retrotransposition that could account for this phenotype. Indeed, offspring raised by LMC mothers with low neurogenesis may possess inherent genetic factors that contribute to the observed decreased in neurogenesis in the DG of pups. This genetic makeup could modulate neurogenesis independently of, or in conjunction with the maternal care received, potentially confounding the observed effects attributed solely to variations in maternal care^68^. Pregnancy induces significant hormonal changes, altering brain structures, particularly in the SVZ^69^. In the SVZ of mice, elevated prolactin boosts neurogenesis early in gestational day (GD) 7, normalizes, then peaks again postnatally, enhancing OB neurons and maternal behaviour^59^. While we did not measure prolactin in our study, suboptimal levels of prolactin in LMC mothers could explain the reduced neurogenesis in the OB, and the associated maternal behaviour. Future studies are needed to assess the role of hormones, including prolactin, progesterone, and testosterone in regulating neurogenesis during pregnancy and postpartum, potentially linking it to natural variation of maternal care and offspring outcomes. Indeed, the drivers of these variations may potentially extend to prenatal factors such as intrauterine position, which affects fetal exposure to hormones like testosterone, estrogen, or prolactin further modulating developmental outcomes and neurogenesis, thus contributing to individual differences in emotional reactivity.

This study highlights that even subtle environmental influences can profoundly shape lifelong emotional trajectories, with adult neurogenesis playing a pivotal role as both a mediator and predictor of these outcomes.

## Acknowledgements

The authors wish to thank, Fulvio Magara and Benjamin Boury-Jamot for help and advice with the behavioural experiments. We would also like to thank the caretakers from the Centre d’Etude Comportementale (CEC). This project was funded by the Swiss National Science Foundation (Grant Number 310030_201015).

## Author contributions

FG, TL, and NT designed research. FG and PvG performed research. FG, TL, PvG, and NT analysed the data. FG, TL, and NT wrote the paper. TL and NT supervised the research. NT obtained the funding.

## Material and methods

### Animals

All experiments were performed on C57Bl/6J male and female mice obtained from Janvier Laboratories (France). After arrival at 7-week-old, mice were left undisturbed for a 1-week-aclimation period. Nulliparous females were used to form breeding pairs, separated from the male after 1 day of mating, and then single-housed for the rest of the experiment. Mice were weighted weekly to track their health status and pregnancy. Mice were randomly distributed into breeding pairs. Mice were maintained under standard housing conditions on corncob litter in a temperature-(23 ± 1°C) and humidity-(40%) controlled animal room with a 12h. light/dark cycle (8h00–20h00), with unlimited access to food and water. All tests were conducted during the light period.

### Preparatory nesting behaviour

Twenty-four hours prior to the nest scoring, the nest material was replaced with 4 fresh squares cotton nestlet, and the mothers remained undisturbed until the next day when scoring was made. On gestational days 14, 15 and 16 at 10am, nests were manually scored as previously described^15^ using an adapted 5 points scale; from 0 to 4. 0: Nesting material is not shredded, no visible nest site; 1: Nesting material is completely or partially shredded, identifiable nest but flat; 2: Nesting material is completely or partially shredded, saucer shaped nest; 3: Nesting material is completely shredded, nest walls have enough height to cover the entire animal, but no closed roof; 4: Nesting material is completely shredded, nest covers the entire animal including a complete roof. When hesitating between two scores 0.5pts were used.

### Anxiety tests

#### Light - Dark test (LDT)

As previously described^70^, the apparatus utilized for the LDT consisted of a white wooden box with two compartments. One was a square compartment without a lid, serving as the light side (40 x 40 cm), while the other was a smaller rectangular compartment with a lid, creating the dark side (20 x 40 cm). These two compartments were connected by a 5 x 5 cm door, and the entire apparatus had a height of 30 cm. The centre of the lit compartment maintained a stable luminosity of 400 lux, while the dark compartment remained without any light source. Mice were introduced into the apparatus in the light side, facing the door, and allowed to explore for a duration of 5 minutes. The mice’s movements were tracked and recorded using the ANY-maze software. In this test, anxiety-like behavior was evaluated based on the time spent in the dark compartment.

#### Open field test (OFT)

The OFT was conducted as previously described^4^ in a rectangular arena (50 × 50 × 40 cm3) illuminated with dimmed light (25 lux). Mice were introduced near the wall of the arena and allowed to explore for 10 min. Analyses were performed using ANY-maze tracking software by drawing a virtual zone (15 × 15 cm2) in the centre of the arena defined as the anxiogenic area. Several parameters were analysed, including the total distance travelled and the time spent in the different zones.

### Anxiety score

The anxiety scores encompassed several anxiety tests to get a general profile of anxiety as previously described^4,5,71,72^. The score is derived from the normalisation of the values for the combination of individual anxiety tests (time spent in the dark chamber during LDT and time spent in thigmotaxis in an OFT). The normalisation involved adjusting each animal’s value by subtracting the minimum value of the entire population and then dividing this result by the difference between the maximum and minimum values of the entire population: (x – min value) / (max value – min value). This method generates scores distributed on a scale from 0 to 1, with a score of 1 indicating high anxiety.

### Maternal behaviour monitoring

Starting on postnatal day (PND) 1, maternal behaviours were systematically recorded twice daily (at 09:00 and 18:00) for two weeks during the light phase, when nursing is most frequent. Using an adapted protocol^12^, cages were observed for 5 seconds every 4 minutes, with each session comprising 10 snapshots per cage, leading to a total of 280 observations per dam over the experiment. Behaviours recorded for each mother were: in the nest: passive nursing, arched-back nursing, contacting pups, licking, or grooming pups, manipulating nest bedding or out of the nest: eating, drinking, self-grooming, rearing. Licking-grooming and arched back nursing are the typical maternal behaviours that serve as the principal criterion in literature to discriminate between high and low maternal care^12,19,20,41^. Hence, the combined percentage of licking-grooming and arched back nursing (LG/ABN) occurrences in both the morning and evening was calculated and correlated to the occurrence in the nest. Dams were classified as high or low maternal care groups based on percentages of occurrence in the nest above or below the cohort mean, which remained consistent across experiments.

### Ultrasonic vocalisation (USV) recording

USVs of each pup upon isolation from the home cage were recorded at PND5 and PND9. Briefly, each individual pup was separated from its mother and littermates and put in a phonically isolated chamber (63×38×42cm) containing a microphone (Avisoft Bioacoustics UltraSoundGate 116H) suspended from the top of the chamber approximately 10 cm from the bottom of the recording chamber. The pup was gently placed in a plastic cup within the recording chamber to minimize movement. Each pup was recorded for 3 min in a range between 10 and 120 kHz. Call detection was provided by an automatic threshold-based algorithm (amplitude threshold: −20 dB; hold time: 5 ms). The accuracy of call detection was verified manually. USVs calls were then analyzed using the oft SASLab Pro software to determine the number of calls, the mean peak frequency, the mean peak amplitude and the mean duration of calls, simple USVs characterized by single, short-duration, frequency-modulated calls and elaborated calls such as ‘complex’, ‘two-components’, ‘composite’, ‘harmonic’ and ‘frequence steps’ that exhibit multiple frequency modulations, varied pitch, and intricate temporal patterns.

### Righting reflex test

To assess the development of the pups’ motor ability, a righting reflex test was performed as already described^73^ on PND5 and PND9. The pup was held in a supine position 5 sec by the experimenter, with all four paws upright. The pup was then released immediately. Righting occurs when the pup flipped over to prone position on the table. The time spent by the pup to flip was used to compute a score: 3: 0-10 seconds to flip, 2: 10-20 seconds to flip, 1: 20-30 seconds to flip and 0: > 30 seconds to flip.

### Cliff avoidance test

The cliff avoidance test was performed as already described^73^ to assess labyrinth reflexes, as well as the strength and coordination ability of the pups on PND5 and 9. The pup was placed at the edge of a flat surface, such that the forepaws and snout of the pup were over the edge. The correct outcome is a protective response, where the pup turns away from the edge of the cliff. Behavior was scored as followed: 0: no movement or falling off the edge, 1: attempts to move away from the cliff but with hanging limbs, 2: successful movement away from the cliff.

### Pups’ retrieval test (PRT)

The pup retrieval test stands out as a predominant test in fundamental and preclinical rodent research for the assessment of maternal behaviour^74^. This test quantifies the mother’s retrieval response to the removal of a pup from the nest. The test was conducted on PND5 and PND9 after 5min of pup’s separation during which USV recording, righting reflex and cliff avoidance tests were performed. Pup retrieval was defined as picking the pup up in the mouth and transporting into the nest. A trial started as the mother was in the nest, and a pup was placed in the most distant corner relative to the nest. The latency to retrieve the pup was measured using a stopwatch. If the pup was not retrieved within 180 seconds, it was placed back into the core of the nest.

### Histology

After all the behavioural experiments the mice were sacrificed with a lethal injection of pentobarbital (10 mL/kg, Sigma-Aldrich, Switzerland) and transcardially perfused with saline solution (NaCl 0.9%) followed by 4% paraformaldehyde (PFA) solution (Sigma-Aldrich, Switzerland). Brains were removed and postfixed with PFA 4% at 4°C overnight. Then brains were transferred in a 30% sucrose solution for 3 days before mounting in OCT compound and stored à frozen at −20°C until slicing. Coronal frozen sections of a thickness of 35 μm were sliced with a cryostat (cryostat, Leica CM3050S) to obtain hippocampal and OB sections conserved in a cryoprotectant solution (30% glycerol, 30% ethylene glycol and 40% PBS 1M) at −20°C until immunofluorescence staining.

### Immunohistochemistry

One out of 6 slices containing hippocampal or olfactory bulb (OB) tissue were chosen to cover the whole dentate gyrus and OB and used for immunostaining. Slices were then incubated 48 h at 4◦C in PBST1% containing 5% of normal goat or donkey serum with the following primary antibodies: rabbit anti-DCX (1:500, Cells Signaling Technology, 4604S), rabbit anti-Ki67 (1:500, Lucerna Chem, ab15580) and mouse anti-Hopx (1:500, Santa Cruz, sc-398703). After 48 hours of incubation, the sections were rinsed 3× 10min in PBST1% and incubated for 3h in either of the following secondary antibodies in PBST1% and 5% of normal goat or donkey serum: goat anti-rabbit Alexa-594 (1:500, Life Technologies, A11037), goat anti-rabbit Alexa-488 (1:500, Invitrogen, A11034), donkey anti-rabbit (1:500, Invitrogen A31573), donkey anti-mouse Alexa-488 (1:250, Invitrogen. A21202). Sections were then rinsed 10min in PBS1x and incubated in 4,6 diamidino-2-phenylindole (DAPI, 1:1000) to reveal nuclei before being rinsed again 3× 10min in PB 0.1M. All images of the immunostained sections of OB and hippocampus were acquired with a Nikon NI-E Spinning disk microscope with a 20X objective. Nikon NIS Elements imaging software was used to do 3D supervised automatic image analysis. The number of labeled cells was counted for each section, in the entire thickness of the section. In the dentate gyrus cells expressing Ki-67 and HopX were counted in an area containing the sub granular zone (SGZ) and the granule cell layer (GCL), whereas the fractional volume occupied by DCX-labeled cells was measured in an area including the sub granular zone, the granule cell layer and the initial part of the molecular layer (ML). In the OB, the fraction volume occupied by DCX-labeled cells was measured in the GCL. The fractional volume is the proportion of DCX^+^ voxels over the total number of voxels, analysed throughout the entire thickness of each section and multiplied by the number of sections analysed.

### Statistical analyses

Statistical analyses were carried out with Prism 9 (GraphPad Software, San Diego, CA 92108, USA), using an alpha level of 0.05. All data are presented as mean ± SEM. Data were tested for normality using the Shapiro-Wilk-test. For normally distributed measures, we used an unpaired, two-tailed Student t-test to estimate differences between the two groups. For no-normally distributed data, we used Mann-Whitney, two-tailed test, whereas when the variances of the groups where not comparable Welch’s test was used. For correlation analyses between either LMC and HMC mother and LMC- and HMC-reared pups at baseline conditions, all possible pairwise correlations were determined by computing Pearson correlation coefficients. Each complete set of correlations was plotted in colour-coded correlation matrices using Prism 9 software. Principal component analyses (PCA) consisted of a series of steps: data selection, scaling, and PCA. For maternal behaviours, we selected seventeen total measures (in the nest, passive nursing, arched-back nursing, contacting pups, licking or grooming pups, manipulating nest bedding, out of the nest, eating, drinking, self-grooming, rearing, resting away from the litter, pups retrieval, time spent in the dark chamber during the LDT, DCX fractional volume in DG, DCX fractional volume in OB and number of pups). For offspring analysis, we selected 18 total measures (DCX fractional volume in the DG, Ki67 positive cells in a mm^3^, time spent in the dark chamber during LDT, anxiety score, total number of calls at PND5, mean duration of calls at PND5, proportion of all simple calls at PND5, proportion of all elaborated calls at PND5, mean peak frequency at PND5, mean peak amplitude at PND5, total number of calls at PND9, mean duration of calls at PND9, proportion of all simple calls at PND9, proportion of all elaborated calls at PND9, mean peak frequency at PND9, mean peak amplitude at PND9 and occurrence in the nest by the mothers). All measures were standardized using a z-score (z = (data point − group mean)/standard deviation) to account for different units across tests. Principal component analysis: We performed PCA using the prcomp() function in Rstudio and calculated the amount of variance explained by each PC using a Scree plot and the loading distribution of principal components using the coefficient outputs from PCA.

