## Supplementary Figures for "Natural variations in maternal behaviour shape trait anxiety and hippocampal neurogenesis in offspring"

### Supplementary Figure 1

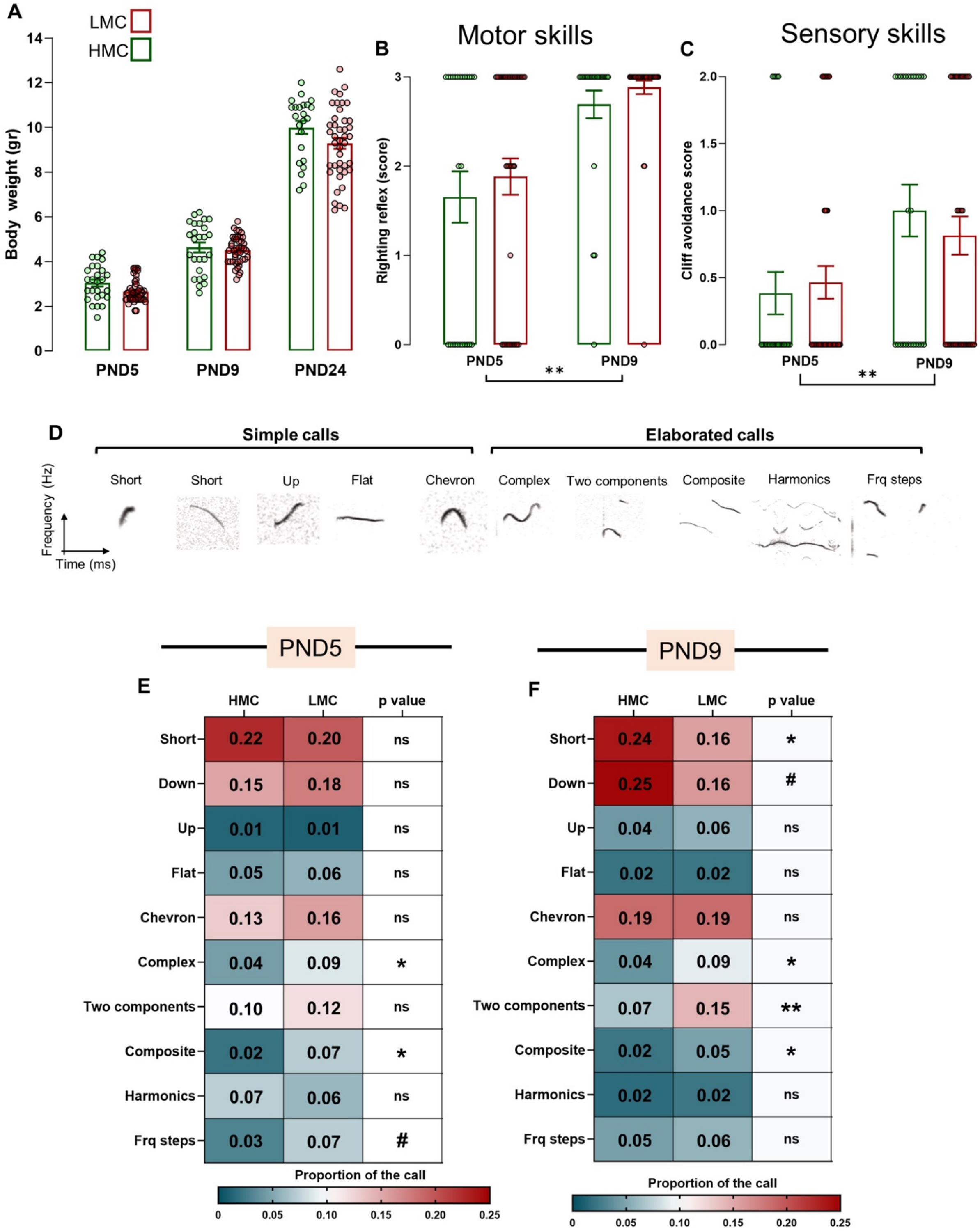

**Supplementary figure 1. Maternal care did not influence other developmental parameters in pups. A.** Body-weight evolution from PND5 to PND24 (time effect: $F_{2,130} = 856.9$, p = <0.001, mixed-effects analysis, n = 26 and 51 per group). **B.** Motor skills at PND5 and 9 analysed through righting reflex (time effect: $F_{1,67} = 32.46$, p = <0.001, Two-way ANOVA, n = 26 and 43 per group). **C.** Proprioception at PND5 and 9 analysed through cliff avoidance score (time effect: $F_{1,67}$, p = 0.006, Two-way ANOVA, n = 26 and 43 per group). **D.** Schematic representation of the type of calls detected and analysed. **E-F.** Heat maps of the proportion of each type of calls analysed, at PND5 (**E.**) and PND9 (**F.**) and the p value for each test used. Histograms show average ± SEM, * p<0.05, ** p<0.01, ***p<0.001, ns = not significant. In **E.** and **F.,** when the data are normally distributed and with comparable variances, unpaired t test is used, whereas for data not normally distributed or with not comparable variances, Welch's test or Mann-Whitney were used.

Supplementary Figure 2

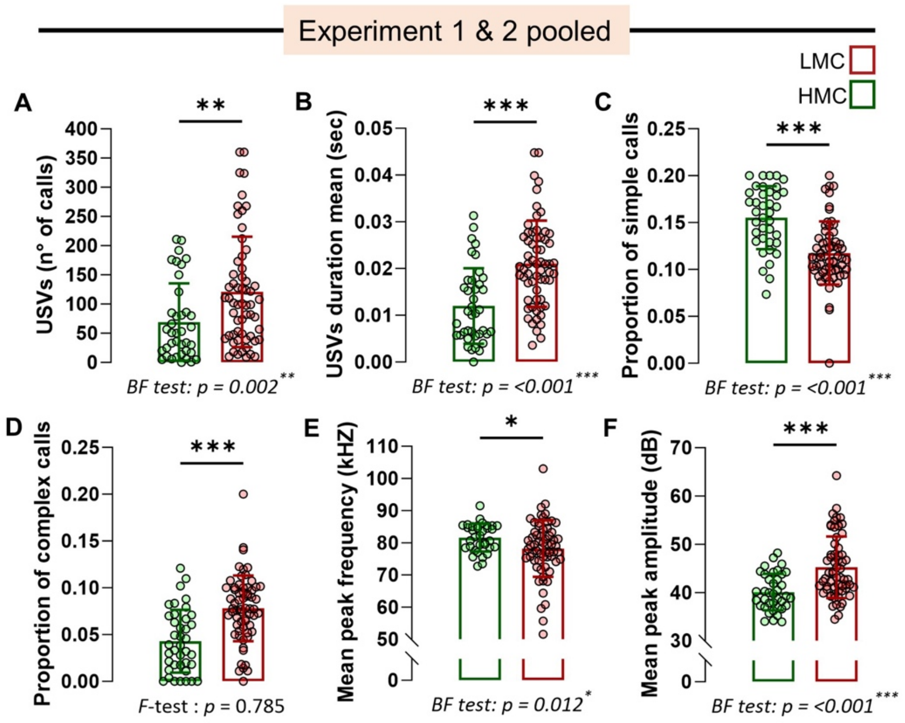


**Supplementary figure 2. Emotional individuality at PND9 is triggered by low maternal care. A.** Total numbers of calls from pups at PND9 pooled from experiments 1 and 2 (Mann-Whitney test, p = 0.002, two-tailed, n = 38 and 59 per group). **B.** Mean duration of calls (Mann-Whitney test, p = <0.001, two-tailed, n = 39 and 62 per group). **C.** Proportion of all simple calls (Mann-Whitney test, p = <0.001, two-tailed, n = 38 and 63 per group). **D.** Proportion of all elaborated calls (t100=4.98, p = <0.001, unpaired t test, two-tailed, n =39 and 63 per group). **E**. Mean peak frequency (Mann-Whitney test, p = 0.038, two-tailed, n = 34 and 62 per group). **F.** Mean peak amplitude (Mann-Whitney test, p = <0.001, two-tailed, n = 38 and 63 per group).

Supplementary Figure 3

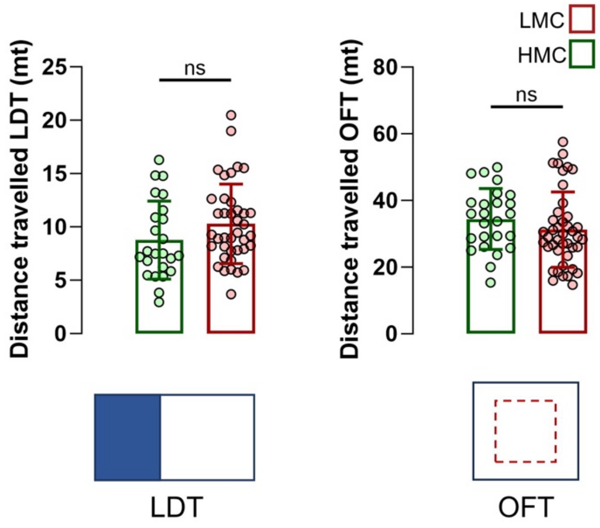


**Supplementary figure 3. Anxiety-related behaviour in juvenile is not due to locomotor deficits. A-B.** Pups raised in HMC and LMC conditions did not show differences in locomotion during LDT ((**A.**) or OFT (**B.**) and PND22-23 (**A.** t61=1.60, p = 0.114, unpaired t test, two-tailed, n = 24 and 39; **B.** t65=1.18, p = 0.241, p = 0.241, unpaired t test, two-tailed, n = 24 and 42).

### Supplementary Figure 4

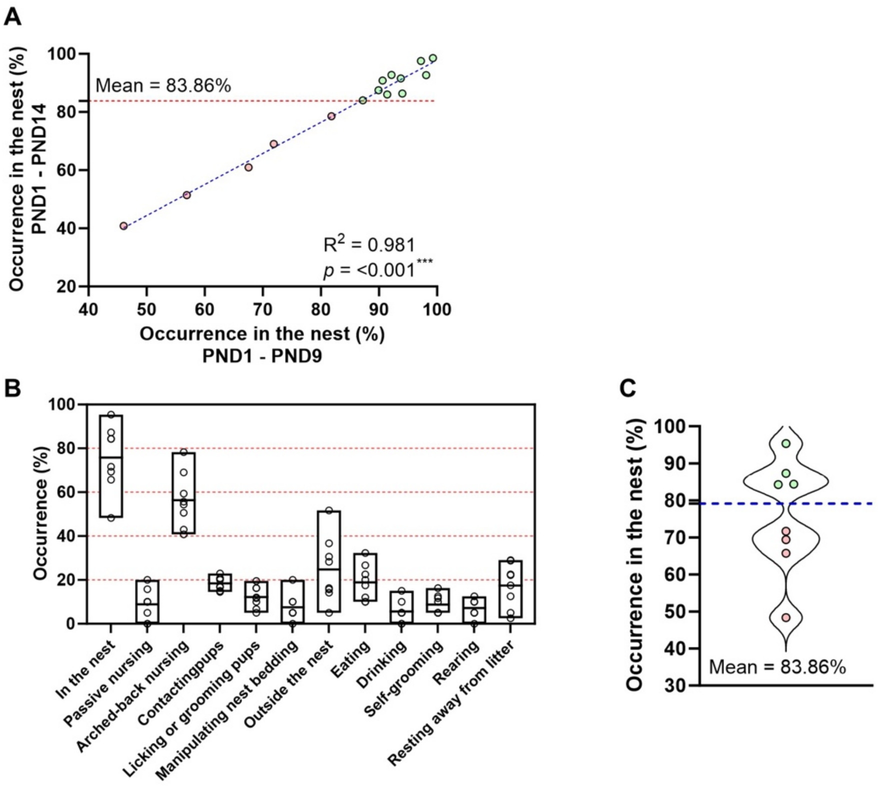


**Supplementary figure 4. Natural variation of maternal care is already observable during the first week of observations. A.** Correlation between occurrence in the nest from PND1 to PND14 and occurrence in the nest from PND1 to PND9 in the first experiment (R2 = 0.981, p = <0.001, simple linear regression). **B.** Representation of the daily observations of maternal behaviours during the experiment 2. **C.** Violin plot showing the distribution and the mean of occurrence in the nest, derived from the observations in the experiment #1, used to discriminate between HMC and LMC.
